# ImmunAL: a frame to identify the immunological markers for Mild Moderated Alzheimer’s Disease applying Multiplex Network Model

**DOI:** 10.1101/2021.10.18.464796

**Authors:** Sagnik Sen, Agneet Chatterjee, Ujjwal Maulik

**Affiliations:** dept. of CSE, Jadvapur University, Kolkata, India

**Keywords:** Immunoprecipitation, Alzheimer’s Disease, Multiplex Network, Autoantibodies

## Abstract

Identification of immunological markers for neurodegenerative diseases resolve issues related to diagnostic and therapeutic. Neuro-specific cells experience disruptive mechanisms in the early stages of disease progression. The autophagy mechanism, guided by the autoantibodies, is one of the prime indicators of neurodegenerative diseases. Identifying autoantibodies can show a new direction. Detecting influential autoantibodies from relational networks viz., co-expression, co-methylation, etc. is a well-studied area. However, none of the studies have considered the functional affinity among the autoantibodies while selecting them from a relational network. In this regard, a two-layered multiplex network based framework has been proposed, whereby the layers consist co-expression and co-semantic scores. The networks have been formed using three distinct cases viz., diseased, controlled, and a combination of both. Subsequently, a random walk with restart mechanism has been applied to identify the influential autoantibodies, where layer switching probability and restart probability are 0.5 and 0.4 respectively. Next, pathway semantic network has been formed considering the autoantibody associated pathways. EPO and IL1RN, associated with a maximum number of pathways, are identified as the two most influential autoantibodies. The network also provides insights into possible molecular mechanisms during the pathogenic progression. Finally, MDPI and CNN3 are also identified as important biomarkers.

**Availability:** The code is available at https://github.com/agneet42/ImmunAL

## I. Introduction

Immunology in neurodegenerative diseases (NDs) have gained a new interest in terms of diagnosis as well as therapeutics [1]. Cell specific immunoprecipitation are associated with functional attenuation which may lead to disease initiation and stochastic progression of the disease [2]. Therefore, autoantibodies (*Aabs*) play a vital role in terms of diagnosis and therapeutics. Computationally modelled effect of immunoprecipitation and neuro-inflammation can provide a clear insight into an early progression of the diseases. However, complicated interplay between differentially co-expressed genes and corresponding co-semantic variation is difficult to observe depending upon singular objective-based networks. In recent years, network models are widely used to fetch influential set of bio-molecules. Besides that, there are many strategies which are implemented on biological data analysis. However, only a single objective has been taken under consideration to establish the relations between pair of nodes within a network architecture, so far. Selecting effective nodes in terms of a single relational objective is biased and is not able to provide comprehensive biological scenarios. The concept of multiplex network can be applied to address the aforementioned issues. Multiplex network is a class of networks where two or more networks share different set of relations among its node. In a single monoplex layer of the multiplex network, the interaction between two nodes are described through edges. Here each of the layers have different mode of interactions. So eventually considering multi-mode interactions from all the networks among homogeneous nodes, the best probable list of genes can be fetched. However, there are two such way which would help to understand the relevance of a node in terms of its presence at different sets of network levels. This is done either by calculating the influence of a node from the network topology or fetching the relatedness between two nodes. Applying both the approaches, it would be an unbiased decision of choosing a set of influential genes where the relatedness between two nodes at different levels can be established. In this article, we have proposed a bi-layered multiplex network where layers are sharing the same proteins with same number of nodes. However, weighted edges, which define the relation between nodes, are different in each layer. In this experiment, the two layers are protein co-expression layer and co-semantic layer. These layers are formed for three different conditions viz., disease case, controlled case and both the cases jointly. In our work, we aim to determine the set of influential proteins from the bi-network by performing a random walk with restart on it to capture the inherent property of the network. Detecting influential nodes in a network has gained prime importance as it controls the nature and depth of information that can percolate in a network. Much work has been done to understand and recognize these seed nodes, which play important roles in understanding the network structure. In [3], the authors evaluate various centrality measures which are used to generate seed nodes and develop a new method based on thresholding the non-seed nodes present in the neighborhoods of the seed nodes. Comin et al. in [4] combine three widely used centralities, betweenness, closeness and eigenvector to show that validated seed nodes have a 95% success rate in scale-free networks. Following the selection process, seed nodes are selected based on their influence in the complex network. As such, many similar models and techniques have been developed for influence maximization. One such model is the Independent Cascade [5] model, where nodes switch between infected, susceptible and inactive stages each of which is associated with a probability of being infected during a particular cycle of spread. Chen et al. [6] describe a community detection based model, where community structures are described out of a network, which narrows down candidate seed selection and finally selects seed nodes based on the purity of the community it belongs to and the position of that node in the network. Genetic algorithm [7] is used to find feasible solutions in the network; whereby a simple 1-point crossover and mutation operator is used. The method assumes no underlying aspects of the network and produces unbiased results. The random walk with restart (RWR) method has been widely used in domains such as epidemic analysis, bioinformatics, video sequencing, autogeneration of captions etc. In [8], researchers use RWR to detect regions in the video sequences which have a temporal salient distribution. They use the transition matrix and restart probability to successfully suppress noise and detect informative features. Jung et al. [9] modify the traditional RWR process to derive influential nodes in a signed network by introducing a signed random walker, whose behaviour modifies accordingly as per the negative/positive direction of the edges. Duc Ha-Le in [10] investigates how the RWR algorithm can be very useful in bioinformatics, especially in ranking related association/interaction problems; RWR has helped in prediction of disease related genes, top k-associations of a protein and to simulate drug target interactions. The next domain of consideration are the multiplex networks, where individual networks are stacked up together; to identify multimodal relationships between a set of nodes. Gilles et al. [11] discover that multiplex networks are better able to capture the community structure in comparison to the aggregated and heterogenous counterparts. They performed their experiments on 4 layers of functional interactions and proved that the multiplex structures allowed for better cohesive in the structure. Multiplex PageRank algorithm has been employed by Li et al. [12] where they solve the complex problem of identifying functional modules in large-scale gene-gene interaction and protein-protein interaction networks.

From the last decade, network algorithms are extremely useful to resolve many biological questionnaires. In [], proteinprotein interaction a network has been analyzed applying the random walk. A complex network representing the pathway membership has been analyzed in [] where three Y2H pathways are combined focusing on 10 pathways information from KEGG. Similarly, a random walk has been applied to predict synthetic lethal biology interactions [13]. In this article, Chipman et al. studied S. cerevisiae and C. elegans with a 95% false-positive rate. Usually, a random walk is applied to understand the affinity between two nodes. However, the starting point is random unless it is pre-defined. Along with that, the algorithm has to start the next iteration from the last visited node. In this regard, two different algorithms have been implemented in this paper for providing comprehensive results. Firstly, eigenvector centrality has been implemented to identify certain nodes which are essentially influential. Subsequently, random walk with restart is another algorithm where we can start again and again from the starting node which also helps to choose another walk with a new sets of nodes. In Koschützk et al. [14], the study shows a comparative analysis between the centralities applied in the biological networks where they have shown five centralities scoring Viz., eccentric centrality, degree centrality, closeness centrality, betweenness centrality and eigenvector centrality for both protein-protein interactions and Transcription Regulation. Based on the correlation among the centrality scoring, eigenvector centrality shows the strongest affinity. Most of the studies where eigenvector centrality has been successfully applied, are complex network problems. Sola Luis et al. [15] shows a mathematical model of eigenvector centrality which is successfully applied on a multiplex network model. The application of the RWR is discussed in Tong et al [16] where they develope model in [16], successfully applied to Corel images and the DBLP database, is faster than other conventional algorithms. The method is successfully applied to Corel images and the DBLP database. Association of diseases and long-non coding RNAs are well known. In [17], a random walk based method has been proposed to predict the association between lncRNAs and diseases. In this case, few major similarity matrices are used to find the probability vector for RWR. The data has been fetched from lncRNA-disease curated database. In recent years, RWR has been implemented on heterogeneous multiplex network. In [18], the new RWR model on multiplex network has been provided. Here, two different monoplex layers have been taken Viz., Protein-Protein interaction network, and co-expression network. The model works comprehensively well. However, none of the models have mapped the functional modifications as well as expression changes. To address such issues in this article we have proposed one framework.

## II. Overview on Random Walk, Random Walk with Restart, and Multiplex Network

This section briefly discussed the basic of random walk, random walk with restart and multiplex network which subsequently use to identify the markers.

### A. Random Walk

A random walk (RW) on a graph is a random movement within a pair of nodes of the graph. The direction of the walk can be decided by a transition probability. Let a monoplex graph be G (V, E), where V denotes a set of vertices and E denotes a set of edges, such that *E_ij_* denotes edges between nodes i and j. Let w(*E_ij_*) denote the weight associated with an edge *E_ij_*. Weighted edges enable quantifying the relationship, thereby allowing varying degrees of relationships throughout a graph. In a RW, the walker moves randomly amongst one of its neighbors at each step. If the walker is at node i, then at the next iteration it selects a node j, from its neighbors. It is a Markov Process predicting or selecting the next step depends only on the present vertex. An N-state Markov Process is therefore characterized by a row stochastic transition matrix *P* of dimension [i,j], where P(i,j) is the probability of moving from node i to node j, where

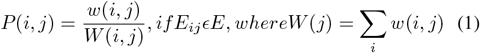

The transition matrix is dependent on the weight of each edge that connects a node i and its neighbors. The transition is guided by higher weighted edges. Hence, if *π^k^* indicates the probability distribution at the k_*th*_ iteration, then,

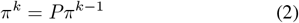

Perron and Frobenius showed when the number of iterations k approaches infinity during a Markov Process, *π* becomes a steady-state of stationary distribution. This distribution is a function of time that a walker spends at this node. This can only be effected by the transition matrix, P. Thus, for the stationary distribution.

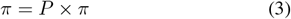

### B. Random Walk with Restart

Random Walk with Restart (RWR) is a special case of RW. It aims at capturing a relationship between a pair of nodes. RWR computes the affinity between two nodes i and j, such that if the walk begins at node i, then the walker chooses a randomly available edge each time. Interestingly, it also has a probability of returning to i, before each iteration. This re-start capability assures that RWR does not get stuck and hence assures the robustness of the method. The steady state distribution of RWR can be formulated as:

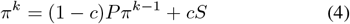

where *π^k^* and *P* are same as Eq. 3, c is the restart probability and S = [s_1_, *s*_2_, *s*_3_,…, *s_N_*]^*T*^, where s_*k*_ denotes the probability of the walk restarting at node k. Each node k∈S, is known as the seed node. These seed nodes are the initialisation points of the random walk. Selection of seed nodes becomes an important decision, because affinity values are generated with respect to these nodes. The restart probability *c* can control the walk. A high restart probability tends to generate walks of shorter length. RWR captures both the global structure of the graph as well as multi-modal relationships between nodes. The latter property has been exploited while developing the random walk with restart on multiplex networks.

### C. Multiplex Network

A monoplex network G is a graph which contains only one layer and only one set of nodes and edges. A multiplex network on other hand, is described by a set of families of graphs G^*m*^ = (V^*m*^,E^*m*^), where m denotes the index of the monoplex networks. A multiplex network is characterized by intra-edges between vertices of the same layer and inter-edges between vertices of different layers. Such kinds of networks allow for modeling of multi-faceted relationships between the vertices. Such inter-edges between two sets of layers are denoted as E^*m*1 *m*2^, which indicates a connection between two graphs G^*m*1^ and G^*m*2^. Each node 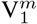, which indicates a node in the *m_th_* monoplex network can have an intra-edge or an inter-edge associated with it. For the purpose of a random walk, each node will have a transition probability associated with it which determines whether the walk will be in the same or between different layer of the network. Fig. 1 explains the structure of a multiplex network.

**Fig. 1.**
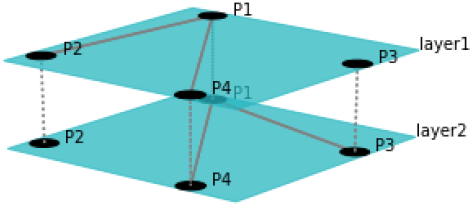
Structure of a multiplex network. Intra-network connections are denoted by a complete line whereas the inter-layer edges are dotted lines. As can be seen, the intra-layer edges are spread across an individual monoplex network whereas the inter-layer edges are connected to only their counterpart nodes across the layers.

## III. Methodology

Mathematical description of the RWR and Multiplex Network have been given in the last section. In this article, we have aimed to apply multiplex network based frame which helps to identify *Aabs*, considering their co-functional similarity. In this case, layer 1 and later 2 are comprised protein co-expression and Go co-semantic respectively. The two layered frame has been discussed below. In the first part, the proposed network architecture has been described. Subsequently, RWR has been discussed in the second phase. The Complete framework has been shown at Fig. 2.

**Fig. 2.**
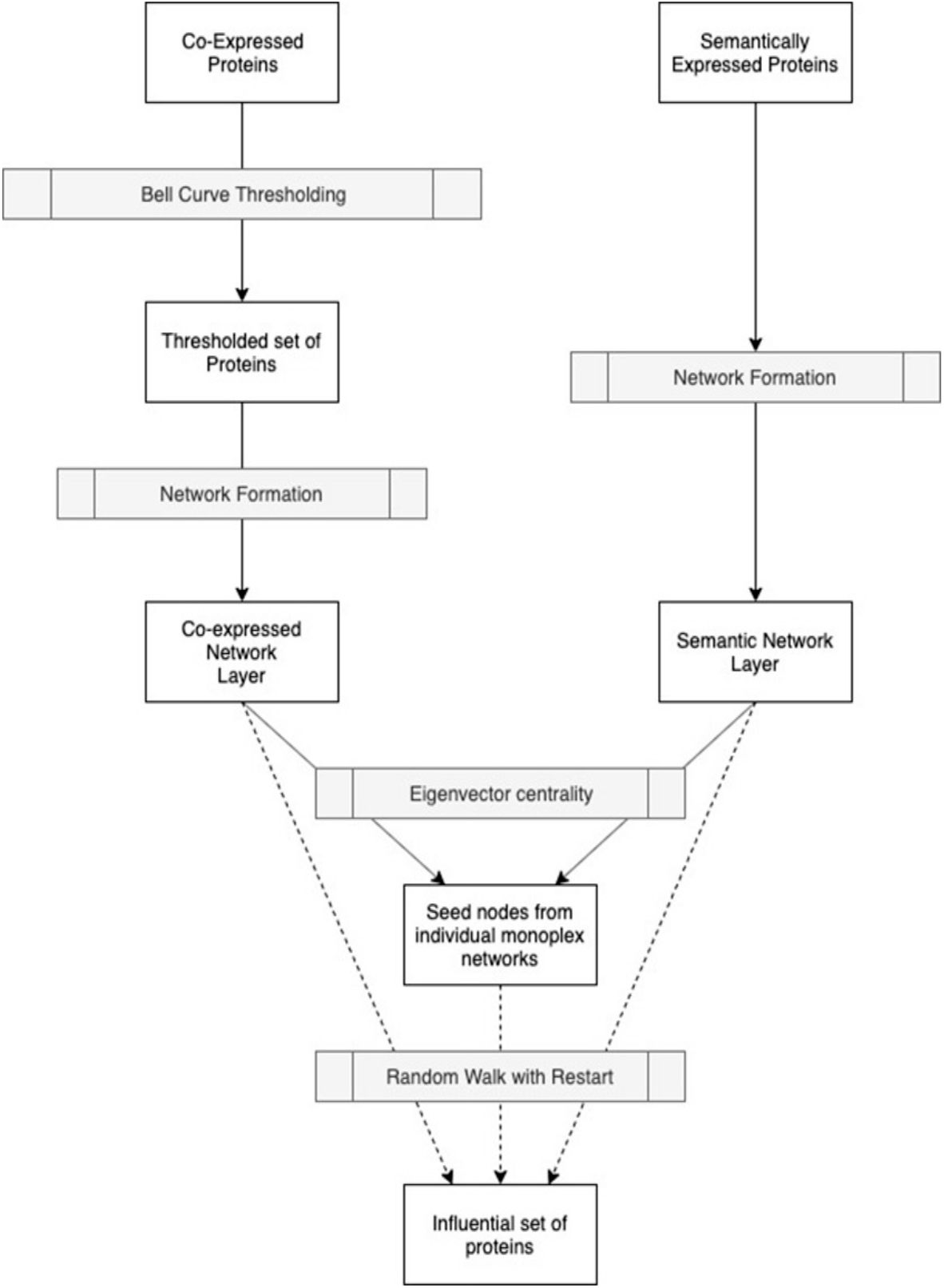
A Schematic Diagram of Proposed Network

### A. Network Sources

The first imperative step in developing the multiplex network is to depict the multi-modal relationships that the nodes share among themself. As mentioned above, each layer in the multiplex layer captures independent relationships between a set of nodes. The following subsections develop a schematic description of each of the monoplex networks:

1. *Co-expressed Layer:* In this article, we have used a protein array of autoantibodies, associated with Alzheimer’s Disease. Initially, a screening has been done to detect differentially functioned protein samples. From the protein array, these chosen samples are expected to be significantly differentiable. In this layer, nodes are representing the *Aabs* (*v*_*c*_1__,…., *v_c_n__* ∈ *V_c_*) whereas weighted edges represent the cooccurrence between them (defined as *e*_*c*_1__,…., *e_c_n__* ∈ *E_c_*) and the weights represent the rate of co-occurrence (from Equ. 5).

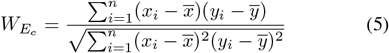

where, *x*_1_,….,*x_n_* ∈ *X* and *y*_1_,…., *y_n_* ∈ *Y* represent the expression values of protein samples. 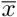 and 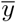 are the population mean. However, only co-expression is not enough for understanding the biological implication of the network. As discussed before, co-expression is showing the rate of co-occurrence of protein *Aabs*. However, it is not necessary that samples are sharing same set of biological processes or pathways. To address the problem, co-semantic layer has been introduced.
2. *Co-semantic Layer:* Like co-expressed layers, co-semantic layer depicts similarity score based on ontology terms Viz., Biological Process, Molecular Function and Cellular Component of the selected protein *Aabs*. Unlike the coexpression layer, this layer provides the information regarding co-existence within a functional system with the rate of cooccurrence. The objective of this layer is to detect whether the samples are sharing the same set of biological process, molecular function and/or cellular component. More elaborately, proteins which are associated with similar kind of biological processes and molecular function at same subcellular localizations, are showing a higher rate of co-semantic score. In this layer, nodes are representing the protein *Aabs* whereas edges are representing the rate of similarity in terms of ontology. The semantic layer is one such layer that can address the aforementioned constraint where the co-occurrence of autoantibodies is allied on association with similar functional systems. To calculate the semantic similarity, we have applied Wang’s method which [19] defined the semantic score of term *Y*. *Y* terms are calculated as *DAG_Y_*=(Y,*X_Y_,E_Y_*), *V_Y_* and *E_Y_* represent set of Go terms and the set of Go terms connecting edges respectively. The *V_Y_* includes term *Y* as well as all its ancestors. Thus, defining the contribution of a GO term *p* to the semantic of GO term *Y* as the *S* – *value* of GO term *p* related to term *Y*. For any of term *p* in *DAG_Y_*, its *S* – *value* related to term *Y, S_Y_*(*p*) is defined as:

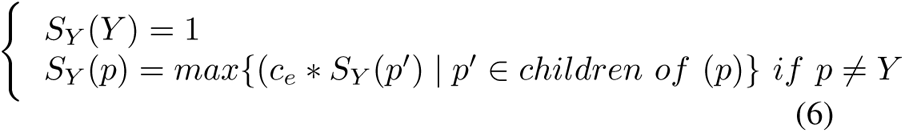 Here *c_e_* is the semantic contribution factor for the edge *e* ∈ *E_Y_* linking GO term *p* with its child term *p*′. After calculating the *S* – *value* for the GO term in *DAG_Y_*, the semantic value of GO term Y, *SV*(*Y*) is defined as:

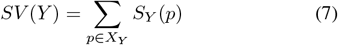 For two given GO term, *Y* and *Z*, the semantic similarity between them is defined as:

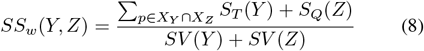 In equation 8, the method proposed by Wang et al. [19] is used to compute the GO semantic similarity (*SS_w_*). Moreover, *S_Y_*(*p*) is the *S* – *value* of GO term *p* related to term *T* and *S_Z_*(*p*) is the *S* – *value* of GO term *p* related to term *Z. X_Z_* is the set of GO terms including term *Z* as well as all its ancestors. Based on the semantic similarity of GO terms, BestMatch Average 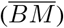 [20] strategy is performed to compute semantic similarity among sets of GO terms associated with the markers associated with a particular pathway, which defined as: 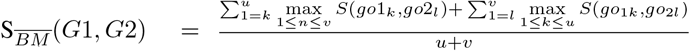 here, gene G1 annotated by GO terms set *GO*_1_ = (*go*_11_, *go*_12_ ⋯ *go*_1*u*_) and G2 annotated by *GO*_2_ = (*go*_21_, *go*_22_ ⋯ *go*_2*v*_).
3. *Bell Curve Threshold:* Due to the constraints of some proteins having no semantic score, it became imperative to perform thresholding to ensure that the kind of proteins in both the layers must be identical. Thus include the proteins in the two layers of the network only after computing their respective semantic scores. The dimensionality reduction is important to ensure that the random walk should not be confined in the correlation network only. The term bell curve [21] is used to define a graphical representation of a normal probability distribution, whose underlying standard deviations from the mean produce a rounded bell form. A standard deviation is a metric used to calculate the variance of dispersion of data in a series of specified values. The “mean” refers to the average of all data points in the data set or sequence.
4. *Network Formation:* The network is formed by developing a complete graph in both the layers, co-expressed and semantic so as to encapsulate all information. If the number of nodes threshold via bell curve is n, then the number of edges in the network are 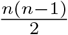.

### B. Influential Nodes selection from Monoplex Network

Selection of seed nodes is an imperative process. For this, we avail the Eigenvector Centrality. A node can be more central if it is in relation with other entities that are themselves central. The centrality of some node does not only depend on the number of its adjacent nodes, but also on their value of centrality. Eigenvector centrality of a node is its summed connections to others weighed by its centrality. The centrality *c* of a node i belonging to an adjacency matrix M is given as:

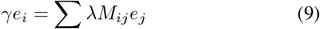

where λ is the associated Eigenvalue of the centrality. In general, the largest Eigenvalue is preferred as the associated Eigenvalue measures the accuracy with which it can reproduce M.

The set of influential nodes from each of the monoplex networks are used as seed nodes for the random walk with restart. These seed nodes are the point of restart for the method, thereby they control the direction of the flow and are critical to the path of the walk that is traversed by the algorithm.

### C. Random Walk with Restart on Multiplex Network

In Random walk with restart on a monoplex network, at each step, the walk can switch to one of its neighbour nodes, or it can restart from the seed nodes. In RWR for multiplex networks, the walk has three possible options, i) it can restart from one of the seed nodes, in its layer, ii) it can continue its intra-network traversal or iii) it can jump to start its internet work traversal. The choice between ii) and iii) is controlled by a parameter μ = 0,…,1; The value of μ indicates that no inter-laying switching happens and it is equivalent to a RWR on a monoplex network. The value of μ can be tweaked to assign importance to different layers, higher values of μ to layer where more traversal is required. If there are two layers *m*_1_ and *m*_2_, and the walker is at node 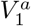, then it can either move to any of its neighbour is layer *m*_1_ or it can move to 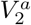. The same is applicable for *m_i_*, where *i* ∈ 1, 2,…, *n*, denoting the number of layers in the multiplex network. In short, it moves to its corresponding node representative in the next multiplex network, whichever is chosen by the parameter μ. Mathematically, as represented in Equ. 10, its equivalent equation would be:

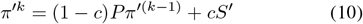

here *π^′k^* and *π^′(k-1)^* are vectors indicating the probability distribution of the particle in the graph. They depict the individual distributions in each layer. The vector S can be defined as μ × *S*_0_. *S*_0_ indicates the initial restart probability whereas = _1,2,…, *n*_, where n denotes the number of layers. *P* is the transition matrix derived out of the adjacency matrix of the multiplex network.

### D. Analysis of the Aabs Using Protein-Protein Interaction and Pathway Semantic Network

The outcomes from the computational frames are highly dependent on the two layered input where functional as well as expression based aspects of the *Aabs*. However, the clear impact of the selected samples can largely be analysed based on their involvement in Protein-Protein Interaction network and pathways. We have applied a two-fold process for the analysis purpose:

1. *Protein Protein interaction network:* From the STRING V11., we have observed the extended interaction partners for the selected proteins based on human species entries. To observe the core interaction module, k-means clustering has been applied. It provides a clear insight into the interaction core.
2. *Pathway Semantic Network and Module Detection:* For the selected sample, a specific list of disease associated pathways are observed. From the literature, the proteins and corresponding pathways can be observed based on their affect on alzheimer’s disease. However, the pathway cross talk can provide newer observation. In this regard, the weighted network *G_path_* has been designed where List of the pathways are considered as nodes ∈ *V_path_* and The semantic similarity between the selected members of the pathways ∈ E_*path*_. The module detection on such weighted network can provide the pathway core. Similarly, the influence of each pathway should be considered during the module detection [22]. The eigen vector centrality based module detection can satisfy such conditions. Therefore, the weighted network is segregated in color modules based on the said algorithm. During the process, eigenvector centrality for each *node* ∈ *V_path_* has also been observed.

## IV. Experimental Results and Discussion

### A. Data Description

In this article, we have used a protein profiling by protein array where the samples are taken from human blood-based serum of Aabs (NCBI ID.-GSE74763) to study the disease progression [23]. There are 50 patient samples from Mild Cognitive Impairment (MCI) stage (Considered as normal for the study) and 50 patient samples from Mild Moderate Alzheimer’s Disease (MMAD) stage. The study has been initiated with 9000 *Aabs* samples.

### B. Experimental Parameters

There are various experimental parameters that determine the traversed network path. The number of influential nodes selected from the monoplex network are 40. The rate of restart is set at 0.3. This allows for the walk to continue at a higher probability. This also lets the traversal scatter across the network, affecting a larger proportion of nodes and not being centered around the seed nodes. The inter-layer switching probability μ is set at 0.5, to allow for equal opportunities for the method to stay in its source layer or move to a different layer. The bell-curve thresholding is done across halves of 2,4,6 and 8 respectively. The networks has been shown in Fig. 3

**Fig. 3.**
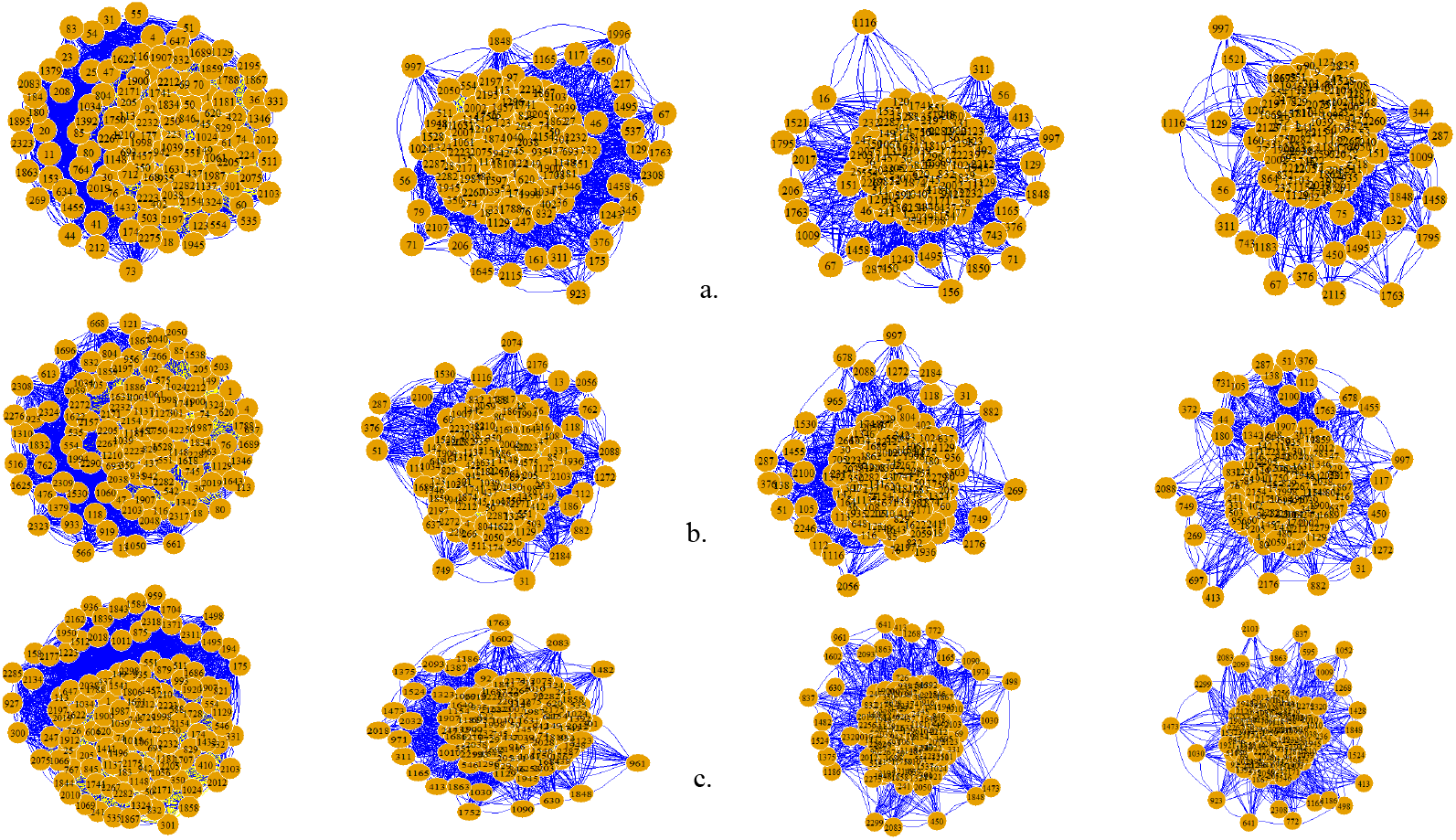
Network diagrams post thresholding of a. controlled and diseases combined cases; b. controlled cases; and c. disease cases

### C. Influential proteins on the Monoplex networks

In the individual monoplex layers, we can identify two different sets of relation, represented in two networks. As mentioned earlier, the two layers i.e., co-expression and co-semantic are homogeneous. However, three individual multiplex networks have been formed. To observe the variability in terms of controlled and disease cases, we have performed the experiment on three samples Viz.,*MultN* – *control, MultN_disease_*, and *MultN_dis-control_*. These results are expected to be more unbiased. As per the description in method section, we have identified top 40 protein samples for each case based on the respective eigenvector scoring. The lists of *Aabs* are given in Supplementary Table 1 - Supplementary Table 4. After thresholding limits as per bell curve distribution, total 12 different sets of influential nodes are identified for each monoplex layers. The experiment has been performed individually for 12 multiplex networks. The influential nodes from each monoplex layer from all 12 multiplex networks are marked as seed nodes. However these 40 influential nodes from each monoplex layer are individually based on the type of objective the layer represents. Therefore, these nodes can be used as starting points for the random walk.

### D. Influential proteins from RWR

The prime objective of using RWR is to identify the influential nodes considering both the layers. As per the basic concept of the RWR, the walk starts from the seed nodes and tries to establish the affinity with other nodes from each layer (here, the number of layers is two). The nodes which have been identified with a high affinity score having association with the maximum number of nodes are known as influential nodes. Such top nodes are detected for all the 12 multiplex networks.

### E. Selecting the Influential Proteins

Now, from the final 12 sets of top proteins sample, a generic set of proteins has been shared by each case shown in Figure 4. Biologically, a shared set of proteins is significantly associated with controlled and disease cases. Also, these proteins are selected even after a stringent thresholding process. Moreover, each of the influential nodes is short-listed depending on the relation objective of each monoplex layer. More elaborately, the selected proteins can be noted as influential in terms of co-expression affinity as well as co-semantic affinity.

**Fig. 4.**
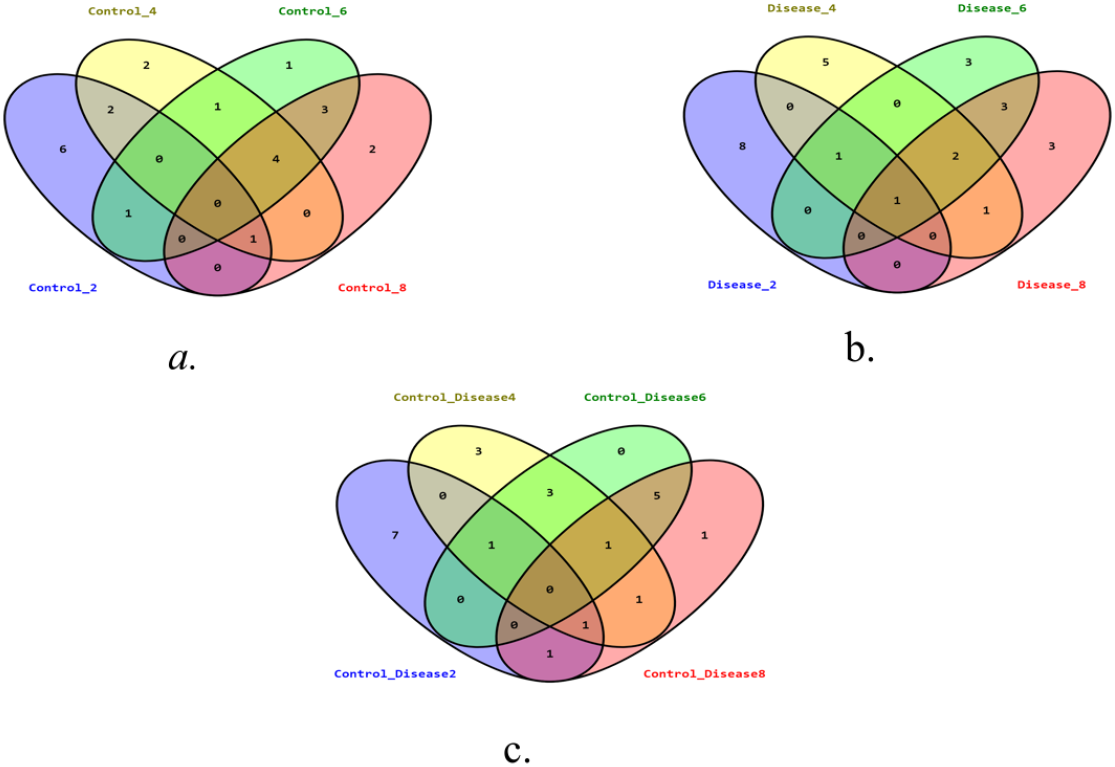
Three Venn Diagrams for three distinctive cases Viz., Controlled, Disease and Controlled-Disease. After segregating from 12 individual sets three Venn Diagram shows specific selected samples.

### F. Analysing the Outcomes using KEGG

We have identified a total of 12 proteins from three separate sets. Five pathways are associated with at least one sample from each set (Controlled, Diseased and Combined). From the EnrichR database [24], we have identified these pathways which are highly involved in the pathogenic progression of AD. In Table II, the list of associated pathways and corresponding protein members are given. From the literature review, all the pathways are directly or partially connected with the pathogenic progression of the AD. In [25], the reduction of NCAM2 has been studied during AD. This reduction has been mediated by amyloid-beta. The reduction of NCAM2 is causing the synapse losses mediated by the synaptic adhesion. Interestingly, cytokine-cytokine receptor pathway is highly associated with pathogenic progression of AD [26]. From the Table I, it has been observed that two prime proteins from the list of RWR disease and combined (IL1RN and EPO) are mediating the aforementioned pathway. Subsequently, a high density of cortisol enhances the risk of dementia [27]. MC2R is noted as one of the leading proteins triggering the cortisol synthesis and secretion pathway.

**TABLE I.**
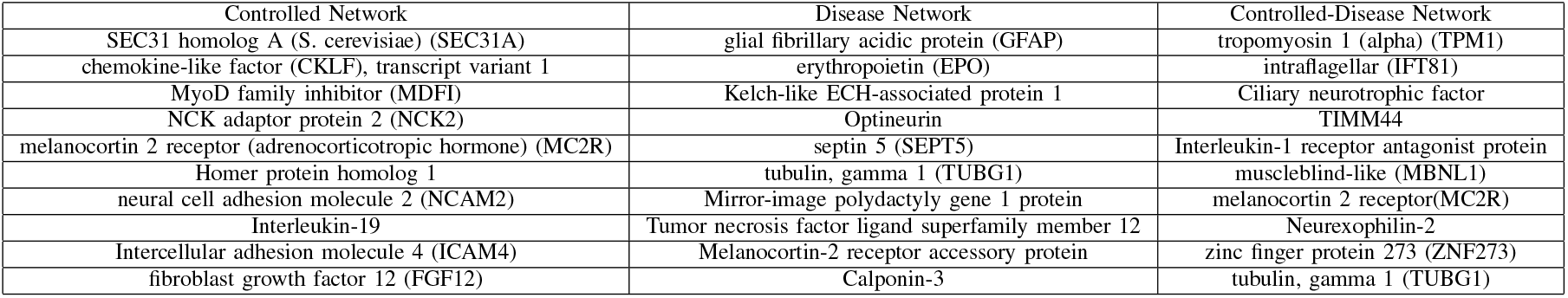
Top 10 proteins selected from each of the networks, when the network is thresholded by a fraction of 4.

**TABLE II.**
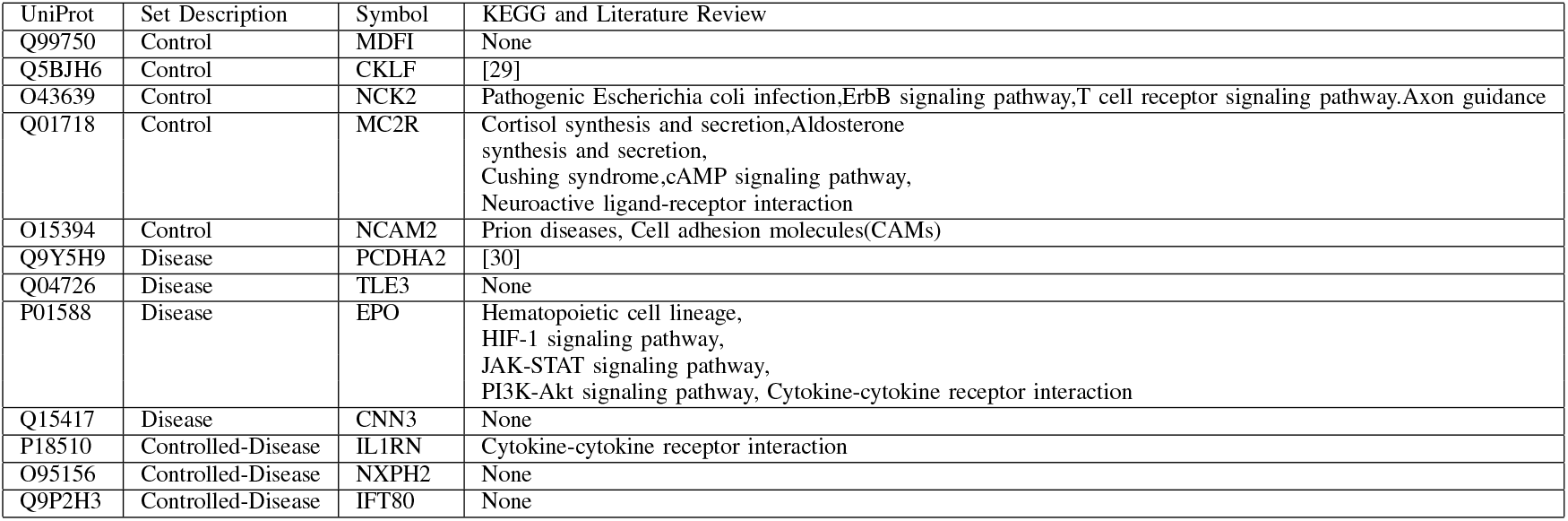
Top 10 proteins selected from each of the networks, when the network is threshold by a fraction of 4.

Similarly, affect of ErbB signaling pathway is largely involved in hippocampus and entorhinal cortex in AD [28]. Interestingly, NCK2 is also one of the members of the ErbB signaling pathway. Prion proteins and AD have long known dependencies. Prion proteins are involved in cell adhesion which has a clear connection with AD pathogenic progression. As mentioned earlier, synaptic cell adhesion is mediated by NCAM2. Here, we have discussed some pathways and their association with AD.

However, other enlisted proteins also have a partial connection with the pathogenic progression. In Figure 5, the given protein-protein interaction network has internal connections among the selected samples. At least one sample from each cluster is from Table II. Interestingly, inter-community connectivity has been governed by the potential nodes. In the network, some samples Viz., NCAM2, PCDHA2, and CKLF have been confirmed with their individual connectivity with the disease. Only two samples i.e., CNN3 and MDPI are the novel findings from the study.

**Fig. 5.**
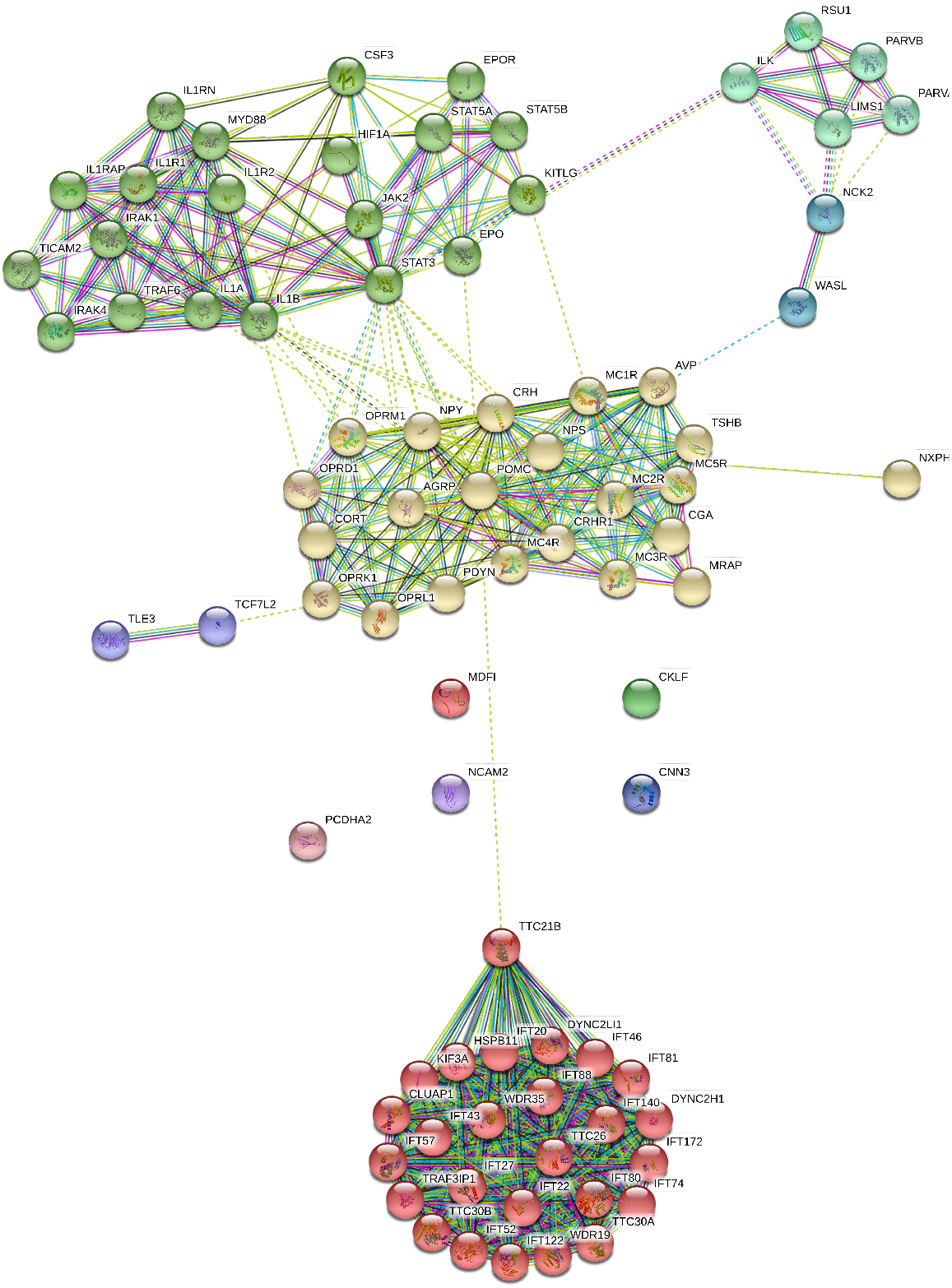
Protein-Protein Interaction Network and Protein Specific Interactive Modules

### G. Analysing Using Pathway Semantic Network

The identified pathway pools were conserved within four markers viz., NCK2, EPO, MC2R, and IL1RN. The pathway crosstalk can be analyzed from pathway semantic network (Shown in Figure 6). In the network, three color modules have four, five and three pathways respectively. Interestingly, each module has at least one pathway which is associated with an immune response e.g., Cytokine-cytokine receptor interaction from cyan module. The module detection algorithm is based on the eigen vector centrality which clarifies the influence of each node in the network. Segregated in the modules, the pathways with maximum eigen vector from each module are Neuroactive ligand-receptor interaction, JAK-STAT signaling pathway, and Axon guidance. However, the average eigen vector centrality of the cyan module is higher than any other module. The members of the respected modules are associated with EPO and IL1RN. Therefore, these two markers can leave an impact on the early immune precipitation which may lead to early stage dementia. Similarly, the pathways with maximum eigen vectors centrality from each module are associated with neuronal activities. Individually, all pathways are contributing in progression of the early stage alzheimer’s disease e.g., the affect of Neuroactive ligand-receptor pathway and cAMP signaling pathway in cognitive responses which lead to advance dementia [31]. However, the understanding can be more stringent by including the essence of cell specificity e.g., the T cell receptor signaling pathway is found as one of the affective pathways in T memory cells as a key regulator for alzheimer’s disease [32]. However, cytokine-cytokine receptor pathway mediated neuroinflammation which is one of the key regulators of alzheimer’s disease, is generic in all the neuronal cells. Finally, EPO and IL1RN are observed as influential biomarkers from the detailed study.

**Fig. 6.**
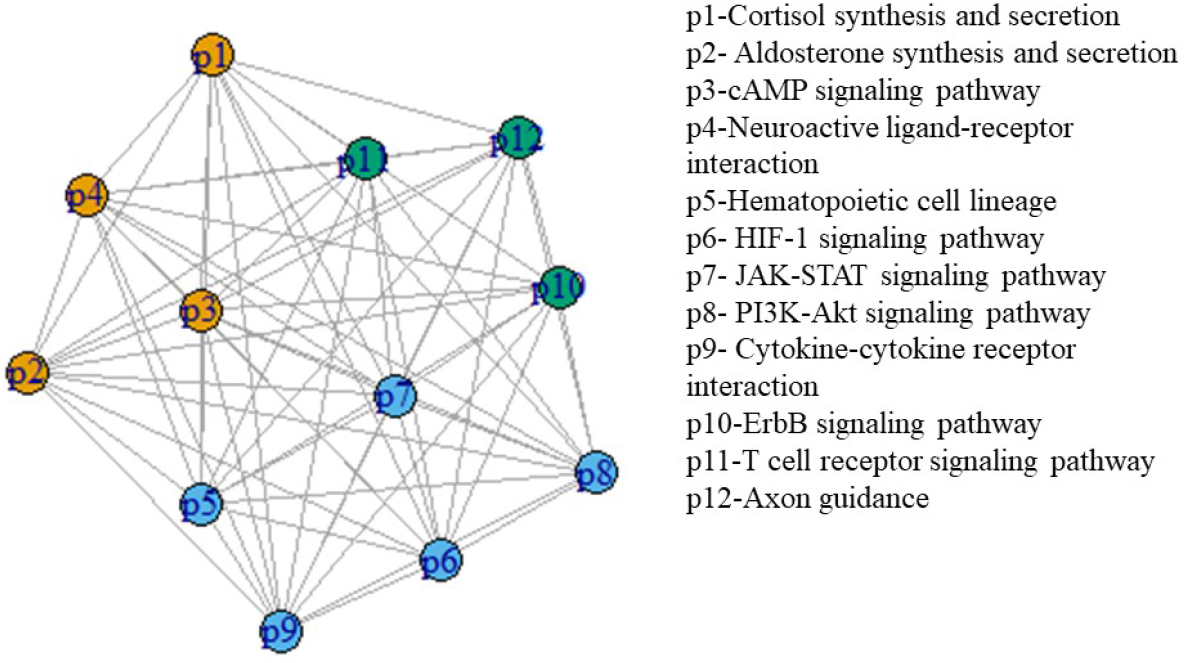
Pathway Semantic Network from identified pathways from Table 2 and eigen vector centrality based three color modules viz., cyan, goldenrod and seagreen.

## V. Conclusion

In this article, the objective was to identify potential *Aabs* markers based on their co-expression and co-semantic patterns. Initially, we have found the 12 indistinctive sets from three cases. As mentioned, samples are taken based on the sharing of GO semantic and expression rate. Therefore, the expression of the potential sample can regulate the maximum number of functional annotations. Interestingly, PPI shows the internal connectivity among the selected antibodies. Also, some of them have literature evidences in early or late onset pathogenic progression. Therefore, the frame has successfully performed to identify the comprehensive markers associated with AD where EPO and IL1RN are identified as influential biomarkers. Finally, we have identified two novel samples i.e., MDPI and CNN3.

## References

[1] R. Hussain, H. Zubair, S. Pursell, and M. Shahab, “Neurodegenerative diseases: regenerative mechanisms and novel therapeutic approaches,” Brain sciences, vol. 8, no. 9, p. 177, 2018.

[2] J. Stephenson, E. Nutma, P. van der Valk, and S. Amor, “Inflammation in cns neurodegenerative diseases,” Immunology, vol. 154, no. 2, pp. 204–219, 2018.

[3] A. Sheikhahmadi, M. A. Nematbakhsh, and A. Shokrollahi, “Improving detection of influential nodes in complex networks,” Physica A: Statistical Mechanics and its Applications, vol. 436, pp. 833–845, 2015.

[4] C. H. Comin and L. da Fontoura Costa, “Identifying the starting point of a spreading process in complex networks,” Physical Review E, vol. 84, no. 5, p. 056105, 2011.

[5] K. Saito, R. Nakano, and M. Kimura, “Prediction of information diffusion probabilities for independent cascade model,” in International conference on knowledge-based and intelligent information and engineering systems. Springer, 2008, pp. 67–75.

[6] Y.-C. Chen, W.-Y. Zhu, W.-C. Peng, W.-C. Lee, and S.-Y. Lee, “Cim: Community-based influence maximization in social networks,” ACM Transactions on Intelligent Systems and Technology (TIST), vol. 5, no. 2, pp. 1–31, 2014.

[7] D. Bucur and G. Iacca, “Influence maximization in social networks with genetic algorithms,” in European conference on the applications of evolutionary computation. Springer, 2016, pp. 379–392.

[8] H. Kim, Y. Kim, J.-Y. Sim, and C.-S. Kim, “Spatiotemporal saliency detection for video sequences based on random walk with restart,” IEEE Transactions on Image Processing, vol. 24, no. 8, pp. 2552–2564, 2015.

[9] J. Jung, W. Jin, L. Sael, and U. Kang, “Personalized ranking in signed networks using signed random walk with restart,” in 2016 IEEE 16th International Conference on Data Mining (ICDM). IEEE, 2016, pp. 973–978.

[10] D.-H. Le, “Random walk with restart: A powerful network propagation algorithm in bioinformatics field,” in 2017 4th NAFOSTED Conference on Information and Computer Science. IEEE, 2017, pp. 242–247.

[11] G. Didier, C. Brun, and A. Baudot, “Identifying communities from multiplex biological networks,” PeerJ, vol. 3, p. e1525, 2015.

[12] J. Li and P. X. Zhao,“Mining functional modules in heterogeneous biological networks using multiplex pagerank approach,” Frontiers in plant science, vol. 7, p. 903, 2016.

[13] K. C. Chipman and A. K. Singh, “Predicting genetic interactions with random walks on biological networks,” BMC bioinformatics, vol. 10, no. 1, pp. 1–11, 2009.

[14] D. Koschützki, K. A. Lehmann, L. Peeters, S. Richter, D. Tenfelde-Podehl, and O. Zlotowski, “Centrality indices,” in Network analysis. Springer, 2005, pp. 16–61.

[15] L. Solá, M. Romance, R. Criado, J. Flores, A. García del Amo, and S. Boccaletti, “Eigenvector centrality of nodes in multiplex networks,” Chaos: An Interdisciplinary Journal of Nonlinear Science, vol. 23, no. 3, p. 033131, 2013.

[16] H. Tong, C. Faloutsos, and J.-Y. Pan, “Random walk with restart: fast solutions and applications,” Knowledge and Information Systems, vol. 14, no. 3, pp. 327–346, 2008.

[17] X. Chen, Z.-H. You, G.-Y. Yan, and D.-W. Gong, “Irwrlda: improved random walk with restart for lncrna-disease association prediction,” Oncotarget, vol. 7, no. 36, p. 57919, 2016.

[18] J. Zhang, Y. Suo, M. Liu, and X. Xu, “Identification of genes related to proliferative diabetic retinopathy through rwr algorithm based on protein-protein interaction network,” Biochimica et Biophysica Acta (BBA)-Molecular Basis of Disease, vol. 1864, no. 6, pp. 2369–2375, 2018.

[19] J. Z. Wang and et al, “A new method to measure the semantic similarity of GO terms,” Bioinformatics, vol. 23, 2007.

[20] C. Pesquita and et al, “Metrics for GO based protein semantic similarity: a systematic evaluation,” BMC Bioinformatics, vol. 9, 2008.

[21] S. Mishra and A. Datta-Gupta, “Chapter 3 - distributions and models thereof,” pp. 31–67, 2018. [Online]. Available: http://www.sciencedirect.com/science/article/pii/B9780128032794000031

[22] A. Dey, S. Sen, and U. Maulik, “Unveiling covid-19-associated organ-specific cell types and cell-specific pathway cascade,” Briefings in bioinformatics, 2020.

[23] C. A. DeMarshall, E. P. Nagele, A. Sarkar, N. K. Acharya, G. Godsey, E. L. Goldwaser, M. Kosciuk, U. Thayasivam, M. Han, B. Belinka et al., “Detection of alzheimer’s disease at mild cognitive impairment and disease progression using autoantibodies as blood-based biomarkers,” Alzheimer’s & Dementia: Diagnosis, Assessment & Disease Monitoring, vol. 3, pp. 51–62, 2016.

[24] M. V. Kuleshov, M. R. Jones, A. D. Rouillard, N. F. Fernandez, Q. Duan, Z. Wang, S. Koplev, S. L. Jenkins, K. M. Jagodnik, A. Lachmann et al., “Enrichr: a comprehensive gene set enrichment analysis web server 2016 update,” Nucleic acids research, vol. 44, no. W1, pp. W90–W97, 2016.

[25] I. Leshchyns’ka, H. T. Liew, C. Shepherd, G. M. Halliday, C. H. Stevens, Y. D. Ke, L. M. Ittner, and V. Sytnyk, “A*β*-dependent reduction of ncam2-mediated synaptic adhesion contributes to synapse loss in alzheimer’s disease,” Nature communications, vol. 6, no. 1, pp. 1–18, 2015.

[26] T. Nagae, K. Araki, Y. Shimoda, L. I. Sue, T. G. Beach, and Y. Konishi, “Cytokines and cytokine receptors involved in the pathogenesis of alzheimer’s disease,” Journal of clinical & cellular immunology, vol. 7, no. 4, 2016.

[27] S. Ouanes and J. Popp, “High cortisol and the risk of dementia and alzheimer’s disease: a review of the literature,” Frontiers in aging neuroscience, vol. 11, p. 43, 2019.

[28] S. A. Meda, M. E. I. Koran, J. R. Pryweller, J. N. Vega, T. A. Thornton-Wells, A. D. N. Initiative et al., “Genetic interactions associated with 12-month atrophy in hippocampus and entorhinal cortex in alzheimer’s disease neuroimaging initiative,” Neurobiology of aging, vol. 34, no. 5, pp. 1518–e9, 2013.

[29] A. R. Zuena, P. Casolini, R. Lattanzi, and D. Maftei, “Chemokines in alzheimer’s disease: new insights into prokineticins, chemokine-like proteins,” Frontiers in pharmacology, vol. 10, p. 622, 2019.

[30] S.-Y. Kim, S. Yasuda, H. Tanaka, K. Yamagata, and H. Kim, “Nonclustered protocadherin,” Cell adhesion & migration, vol. 5, no. 2, pp. 97–105, 2011.

[31] J.-f. Liu, A.-n. Hu, J.-f. Zan, P. Wang, Q.-y. You, and A.-h. Tan, “Network pharmacology deciphering mechanisms of volatiles of wendan granule for the treatment of alzheimer’s disease,” Evidence-Based Complementary and Alternative Medicine, vol. 2019, 2019.

[32] D. Gate, N. Saligrama, O. Leventhal, A. C. Yang, M. S. Unger, J. Middeldorp, K. Chen, B. Lehallier, D. Channappa, B. Mark et al., “Clonally expanded cd8 t cells patrol the cerebrospinal fluid in alzheimer’s disease,” Nature, vol. 577, no. 7790, pp. 399–404, 2020.

